# Enhancing single-cell cellular state inference by incorporating molecular network features

**DOI:** 10.1101/699959

**Authors:** Ji Dong, Peijie Zhou, Yichong Wu, Wendong Wang, Yidong Chen, Xin Zhou, Haoling Xie, Yuan Gao, Jiansen Lu, Jingwei Yang, Xiannian Zhang, Lu Wen, Wei Fu, Tiejun Li, Fuchou Tang

**Author notes:** Correspondence (W.F.), (T.L.), (F.T.). These authors contributed equally (Ji Dong, Peijie Zhou, Yichong Wu, Wendong Wang, Yidong Chen, Xin Zhou).

## Abstract

In biological systems, genes function in conjunction rather than in isolation. However, traditional single-cell RNA-seq (scRNA-seq) analyses heavily rely on the transcriptional similarity of individual genes, ignoring the inherent gene-gene interactions. Here, we present SCORE, a network-based method, which incorporates the validated molecular network features to infer cellular states. Using real scRNA-seq datasets, SCORE outperforms existing methods in accuracy, robustness, scalability, data integration and removal of batch effect. When applying SCORE to a newly generated human ileal scRNA-seq dataset, we identified several novel stem/progenitor clusters, including a Cripto-1+ cluster. Moreover, two distinct groups of goblet cells were identified and only one of them tended to secrete mucus. Besides, we found that the recently identified *BEST4+OTOP2+* microfold cells also highly expressed *CFTR,* which is different from their colonic counterparts. In summary, SCORE enhances cellular state inference by simulating the dynamic changes of molecular networks, providing more biological insights beyond statistical interpretations.

## Introduction

Instead of executing function in isolation, genes tend to form complex molecular networks and function in conjunction to determine the cellular or organismal phenotypes^1, 2^. Although it has been recognized for a long time that molecular networks are dynamic during organismal development and differentiation, how to simulate this process is still challenging.

With the rapid development of single-cell RNA sequencing (scRNA-seq) techniques, nowadays, researchers can easily profile the transcriptome of large number of cells at single-cell resolution, and identify new cell types and intermediate cellular states within a certain organismal system^3–5^. To fully utilize these rich datasets, many efficient computational methods have been developed, such as Seurat, SCANPY, and SINCERA^6–8^ However, these methods usually calculate the transcriptional similarity inferred from individual genes to do cell clustering analysis and detect the marker genes that can uniquely distinguish the identified cell types, while ignore the integrated and synergistic nature of molecular interaction networks, which in fact formulate the specific cell fates and provide more biological insights. In addition, the performance of individual gene-based methods can be influenced by the prevalent dropout events and other experimental artifacts in single-cell sequencing data, and the performance strongly depends on the subjective selection of highly variable genes in the pre-processing steps.

Recently, several methods such as SCENIC, PAGODA, and CSN are proposed to identify cell types by incorporating some biological knowledge-based information, which utilizes transcription factor-based regulatory networks, functional modules, and cell type-specific networks to facilitate the biological interpretation, respectively^9–11^. However, due to strong technical noises there are still no optimal methods to accurately infer the gene-gene or cell-cell relationship from the sparse scRNA-seq datasets^12^. In the meantime, accumulating public data on molecular interactions derived from sound experimental or computational evidences can provide rich prior biological knowledge to reduce the false-positive rate of molecular network inference ^13, 14^ For example, with the aid of protein-protein interaction (PPI) databases, SCENT is able to infer the single-cell entropy, while netSmooth can temper the noisy scRNA-seq data^15, 16^.

We hypothesize that during the organismal development and differentiation, the transition from a cellular state to another is accompanied by the destruction of critical molecular networks of the former cellerular state and the reconstruction of novel ones (Fig. 1a). In view of this, we introduced a new method, **SCORE** (**S**ingle-**C**ell m**O**lecula**R** n**E**twork), to simulate the dynamic changes of molecular networks from scRNA-seq datasets by incorporating the experimentally validated and high-confidence molecular interaction information from public databases. We validated the accuracy, robustness and scalability of SCORE to uncover cell states using gold-standard scRNA-seq datasets. The performance demostrated the superiority of the proposed method over previous proposals. Finally, with SCORE, we accurately integrated five human fetal datasets and analyzed a newly generated human adult ileal epithelium dataset. SCORE is freely available in https://github.com/wycwycpku/RSCORE.

**Fig. 1.**
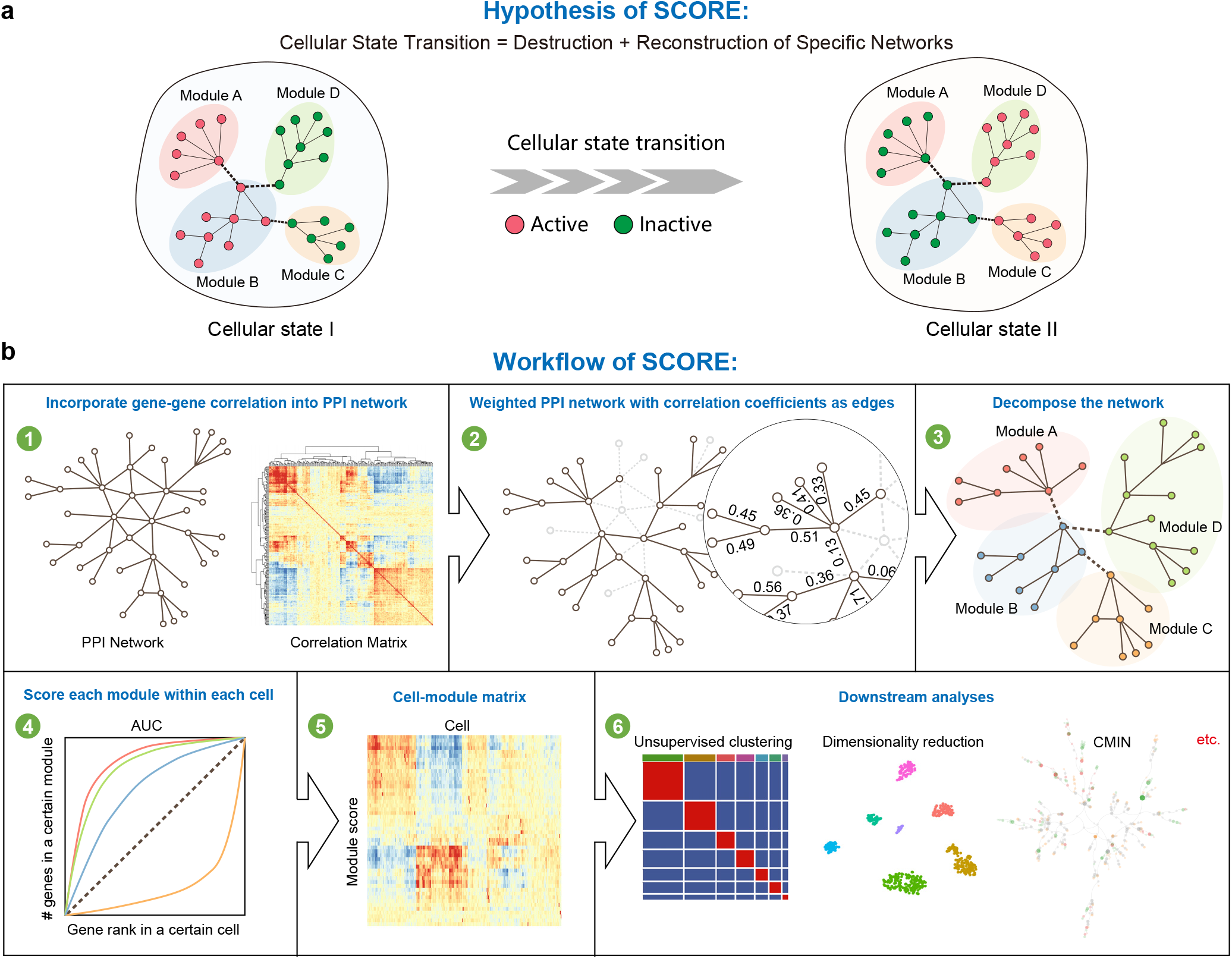
The workflow of SCORE. **a,** SCORE assumes that the cellular state transition is associated with the activation/inactivation of functional modules of molecular interaction network, which can be inferred from single-cell transcriptome data. **b,** In the SCORE workflow, the input protein-protein interaction (PPI) network from public database and gene correlation inferred from single-cell dataset are trimmed to construct a weighted molecular interaction network (WMIN). The random walk approach is then applied to WMIN to decompose molecular interaction modules via a consensus strategy, and the activation score of modules for each cell is calculated from AUCell. Downstream analysis are performed based on the obtained cell-module activity matrix to cluster and visualize the cells against technical variations, and construct the characteristic molecular interaction network (CMIN) for each cellular state.

## Results

### 1. The workflow of SCORE

In brief, as we reason that genes execute functions through interacting with other genes in molecular networks, we need to extract the molecular networks that undergo dynamic changes among different cellular states (Fig. 1b). Thus, we first generate a weighted gene-gene Pearson correlation network. For scRNA-seq datasets, Pearson correlation coefficient can cover more authentic gene-gene relationships compared with other algorithms, although it will also introduce many false positives. To reduce the false positive rates, we use a curated PPI network to trim the gene-gene relationships; likewise, data-irrelevant interactions of PPI network are pruned by the weighted correlation network. The obtained weighted network is then decomposed into numbers of small molecular networks, termed as modules, by random walk algorithm. Finally, each module is scored within each cell using the AUCell algorithm^9^, and a cell-module matrix is obtained, which represents the activity of individual module within each cell.

Next, this cell-module matrix can be utilized to perform downstream analyses such as visualization, clustering, and cell lineage analysis. Importantly, inspired by the concept of Steiner Tree in graph theory, SCORE also constructs the characteristic molecular interaction network (CMIN) to annotate a certain cellular state (see Methods for more details). For the convenience of users, SCORE is seamlessly compatible with the popular R package Seurat^6^.

### 2. Evaluation of the accuracy, robustness, and efficiency of SCORE

To verify the rationale and assess the performance of the algorithm, we applied **SCORE** to a gold-standard scRNA-seq dataset for cell clustering analysis, with 561 cells derived from seven human cell lines (GSE81861) after quality control^17^. Two experimental batches exist in the cell lines for GM12878 lymphoblastoid cells and H1 embryonic stem cells. In the previous literature, several clustering methods were benchmarked without providing the additional information on batch identity, and the reference component analysis (RCA) achieved the highest adjusted rand index (ARI) of 0.91, while other methods showed inferior performance (All=HC: 0.66; HiLoadG-HC: 0.53; BackSPIN: 0.64; RaceID2: 0.15; Seurat: 0.70) (Fig. 2a). Here, we intend to continue the benchmarking in the same set-up, by comparing the performance of SCORE with other state-of-art feature extraction methods based on gene regulatory network, and highlight the strength of SCORE in the accurate identification of the cell lines and the effective alleviation of undesired experimental noises.

**Fig. 2.**
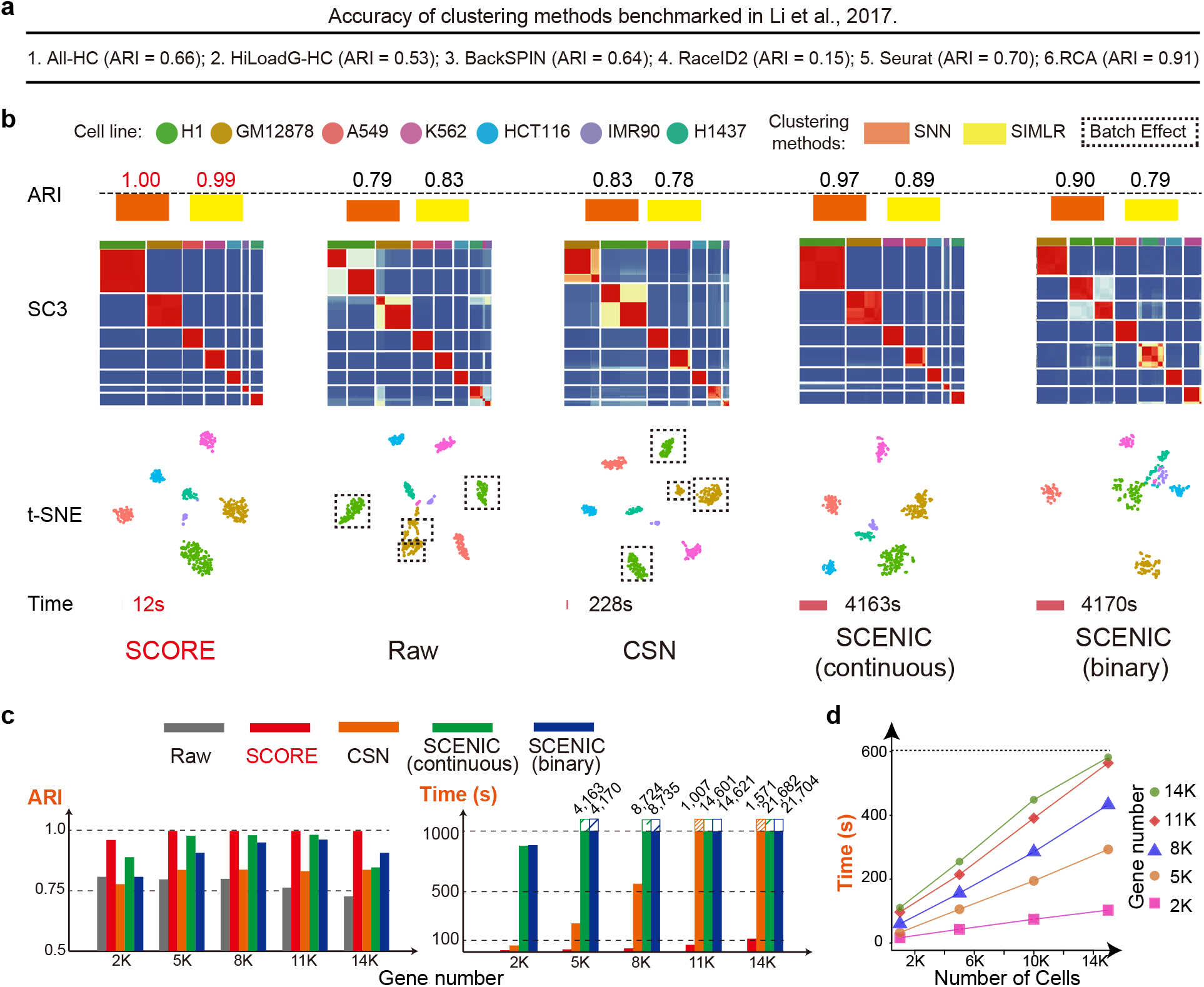
The performance assessment of SCORE. **a,** Accuracy of clustering methods evaluated on the gold-standard cell-line benchmarking dataset in Li, et al. ^17^ **b,** The performance of SCORE on the cell-line benchmarking dataset was compared with the direct analysis on raw expression matrix within Seurat pipeline (denoted as Raw) and other gene regulatory network (GRN) based methods (i.e. CSN and SCENIC with continuous/binary features). SCORE outperforms other methods in terms of both clustering accuracy (indicated by Adjusted Rand Index: ARI of different clustering approach SNN and SIMLR, as well as the batch removal effect revealed by the SC3 similarity matrix and t-SNE plot) and running time. The number of input highly variable genes is set as 5,000, and the colors in t-SNE plot denote cell line identities collected in the experiments. **c,** The assessment of SCORE robustness to feature selection in gold-standard dataset and comparison with other methods. SCORE always achieves the highest ARI and shortest running time regardless of the number of selected HVGs. **d,** The scalability of SCORE with the increase of numbers of input genes and cells, tested by down sampling of the integrated human fetal datasets. The running time of SCORE scales almost linearly with the number of cells, and does not witness the significant increases as the gene number exceeds 11,000. The implementation of SCORE completes within 10 minutes with 15,000 cells and 14,000 genes.

As shown in Fig. 2b, SCORE yielded the highest accuracy, with the ARI amounting to 1.00 (clustering by SNN^18^) and 0.99 (clustering by SIMLR^19^), respectively. Both the t-Distributed Stochastic Neighbor Embedding (t-SNE) plot and SC3^20^ similarity matrix demonstrated that the batch discrepancy was removed for the same cell lines, while the sharp distinction among different cell lines was still retained. In comparison, the direct clustering based on expression matrix within Seurat pipeline (Raw) confronted with limitations, by blurring the distinct cell identities (e.g. K352 and H1437 or IMR90 cells) and discriminating the unwanted experimental batches in both GM12878 and H1 cell lines (indicated by the separate clusters in t-SNE plot and independent blocks in SC3 matrix in Fig. 2b). Despite that the cell-specific network (CSN) method separated different cell lines, clustering variations induced by technical artifacts seemed to be strengthened. Clustering guided by SCENIC continuous features generated relatively satisfactory results comparable to (but still inferior than) the analysis by SCORE, but with considerable more computational costs.

In addition, the performance of SCORE was robust under the selection of input variable genes for analysis. As shown in Fig. 2c and Supplementary Figs. 1-4, SCORE always outperformed other methods for all the selected gene numbers ranging from 2,000 to 14,000, both in terms of clustering ARI and the removal of experimental batch effects, as indicated by the SC3 similarity matrices and t-SNE plots. As long as the gene number exceeded 2,000, SCORE could robustly recover the cell line identities with the optimal resolution. In comparison, while SCENIC achieved satisfactory clustering results with 5,000-11,000 genes, the clustering ARI was significantly reduced for the case of 14,000 genes due to the severe batch effects.

The implementation of SCORE is also efficient, mainly due to the fast module decomposition of PPI by random walk distance, and the highly parallelizable calculation of module activity matrix. With 14,000 top variable genes, SCORE finalized the analysis within 2 minutes, while the implementation of SCENIC lasted for 6 hours in R. In fact, from the running time tests over various groups of genes (Fig. 2c), the implementation of SCORE was typically 10-20 times faster than CSN and 100-400 times faster than SCENIC. Notably, SCORE could handle the task of 15,000 cells and 14,000 genes in less than 10 minutes (Fig. 2d).

Overcall, application of SCORE to the gold standard dataset validates our rationale- the incorporation of molecular interaction network in scRNA-seq analysis can facilitate the accurate cell identity dissection and reduce the technical artifacts in experiments. The comparisons with other methods also suggest the capability and potential of SCORE for the unbiased and efficient analysis, and integration of large-scale transcriptome datasets, which will be explored below.

### 3. Integration and multi-scale comparison of five human fetal datasets

Using scRNA-seq techniques, many studies have uncovered unprecedented findings in terms of human fetal development. However, most of these studies focused on just one organ or tissue and failed to consider the human fetal development in a holistic view. In addition, it is also a challenge for existing methods to integrate the datasets sampled from different organs and different time points because of batch effects introduced by varied dissociation protocols, organ-specific differences, etc. Given this, we used SCORE to integrate five high-quality human fetal datasets previously generated by our group, including fetal gonads (overy and testis)^21^, heart^22^, kidney^23^, prefrontal cortex (PFC)^24^, and cerebral cortex^25^, spanning from 4 to 26 weeks of fetal development. Importantly, we re-organized the five datasets using uniform pipeline and format, which provided a rich and convenient resource for studying human fetal development (https://github.com/zorrodong/HECA). To evaluate the integration result, we utilized the mixability of common cell types shared by these datasets, such as immune cells, erythroid cells, and endothelial cells across different organs. Notably, SCORE obtained a reasonable result with cell types grouped by their own identities, while Seurat pipeline on individual gene expression matrix (Raw) failed to cluster these common cell types (Fig. 3a,b and Supplementary Fig. 5). We also used two batch effect correction methods, Harmony^26^ and CCA^27^, to integrate these datasets; however, they all exhibited defective results. Even worse, both Harmony and CCA introduced serious artificial results and unrelated cells were wrongly grouped together.

**Fig. 3.**
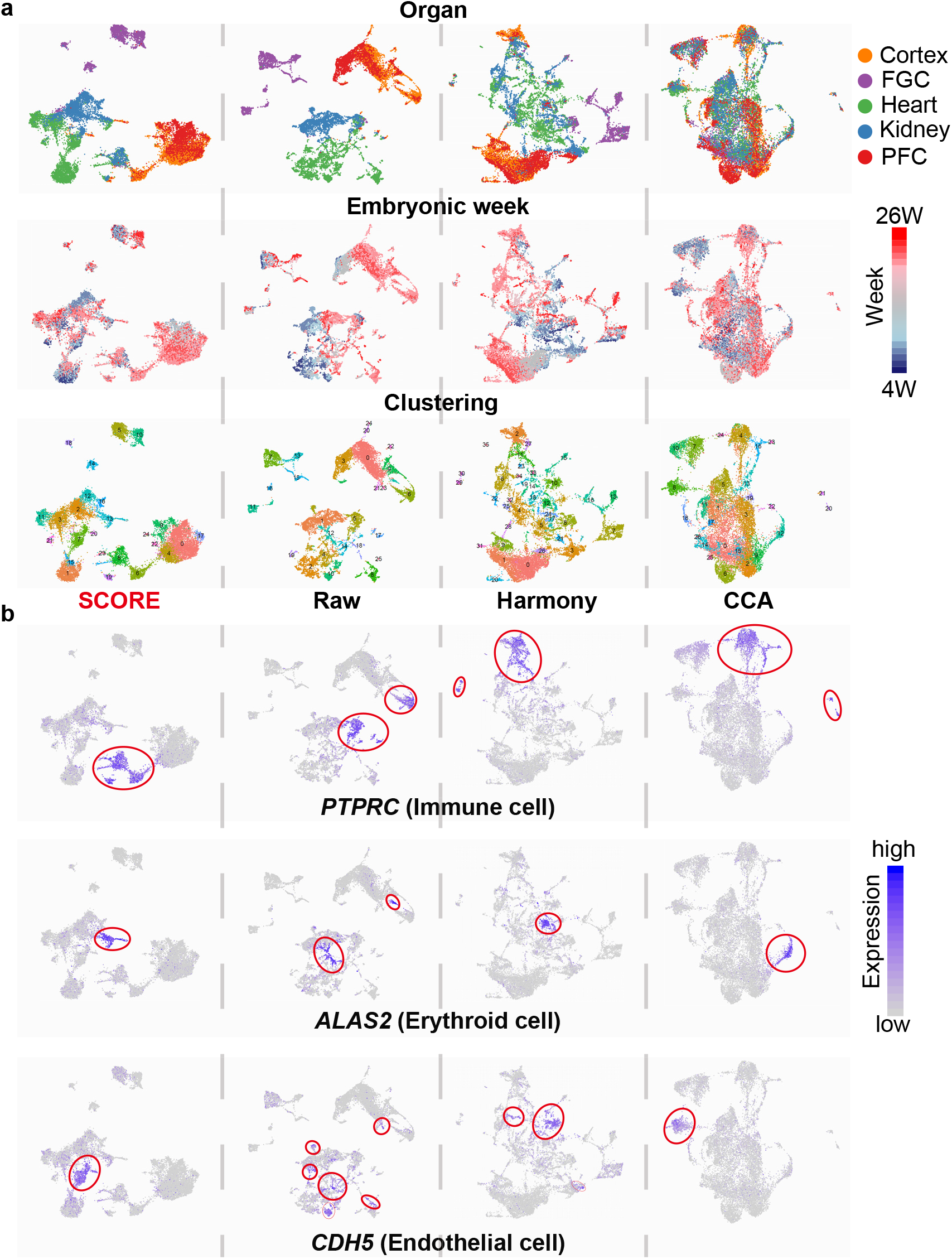
Unbiased integration of five human fetal datasets. **a,** The UMAP visualization of different data integration results implemented by SCORE, direct merge of datasets (Raw), Harmony and CCA (aligned from left to right in each row). The cells are colored by organ information (the top row), developmental week information (middle), and clustering results based on integrated data (bottom), respectively. **b,** The UMAP visualization of data integration results by SCORE and other methods, with cells colored by the expression level of marker genes for immune (top), erythroid (middle), and endothelial (bottom) cells. The cells that highly express the marker genes are denoted by the red circles.

In addition, we also analyzed each of these five datasets separately (Supplementary Figs. 6-10). Take the human fetal kidney dataset as an example. Standard clustering using Seurat pipeline showed a chaos state, especially in renal interstitium (RI) (Supplementary Fig. 6a,b). Specifically, there was a distinctly isolated cell population of 19-week fetal renal cells, which might be caused by batch effects. In contrast, SCORE presented a much better result. All fetal cells were first separated into renal cell populations and non-renal cell populations, including immune cells (IMMs), erythrocytes (ERs), and endothelial cells (EDs). Renal cells comprising glomerular cells, renal capsule cells, and renal tubular cells were then classified into 9 clusters according to the anatomic structure of nephron, namely, cap mesenchyme (CM), podocytes (PDs), proximal tubule (PT), loop of Henle (LH), distal convoluted tubule (DT), collecting duct (CD), intraglomerular mesangium (MG), extraglomerular mesangium (EM), and renal interstitium (RI). Importantly, the UMAP plot using SCORE displayed an accurate developmental trajectory of nephrons from the cap mesenchyme to the formation of epithelial tubules (Supplementary Fig. 6c-e). For other four datasets, SCORE also performed better than Seurat. In heart dataset, SCORE firstly divided cardiomyocyte cells (CMCs) into compact CMCs and trabecular CMs, and then portioned compact CMCs into atrial CMCs (CMCs-A) and ventricular CMCs (CMCs-V), which was consistent with the anatomic structures of heart (Supplementary Fig. 7). In two cortex datasets and gonad dataset, slight batch effects were detected in Seurat results but not in SCORE results (Supplementary Figs. 8-10).

### 4. Expression landscape of human adult ileal epithelium

Although small intestine epithelium has been studied widely in murine, a comprehensive expression landscape in human is still lacking. To explore the cellular diversity of human small intestine epithelium, we sampled the ileal crypts from two patients suffered from right-sided colon cancer while their ilea were relatively normal (Supplementary Fig. 11a). scRNA-seq libraries were generated using the 10X v3 Kit, which could guarantee both throughput and quality (Supplementary Fig. 11b). Compared with fetal datasets, adult ones tend to be more complex due to more individual differences. As expected, Seurat pipeline grouped cells according to the patient individuals, which indicated the existence of batch effect and would result in confusion and inaccuracy for downstream analyses (Fig. 4a). In contrast, SCORE successfully eliminated these unwanted effects and identified 19 clusters, which covered all the known cell types, namely, stem/progenitor cells, paneth cells, tuft cells, goblet cells, enteroendocrine cells, microfold (M) cells, enterocytes (Fig. 4b and Supplementary Table1). The differentially expressed genes (DEGs), differentially activated modules (DAMs) and the inferred differentiation potency all supported the clustering accuracy of SCORE (Fig. 4c,d and Supplementary Fig. 12a).

**Fig. 4.**
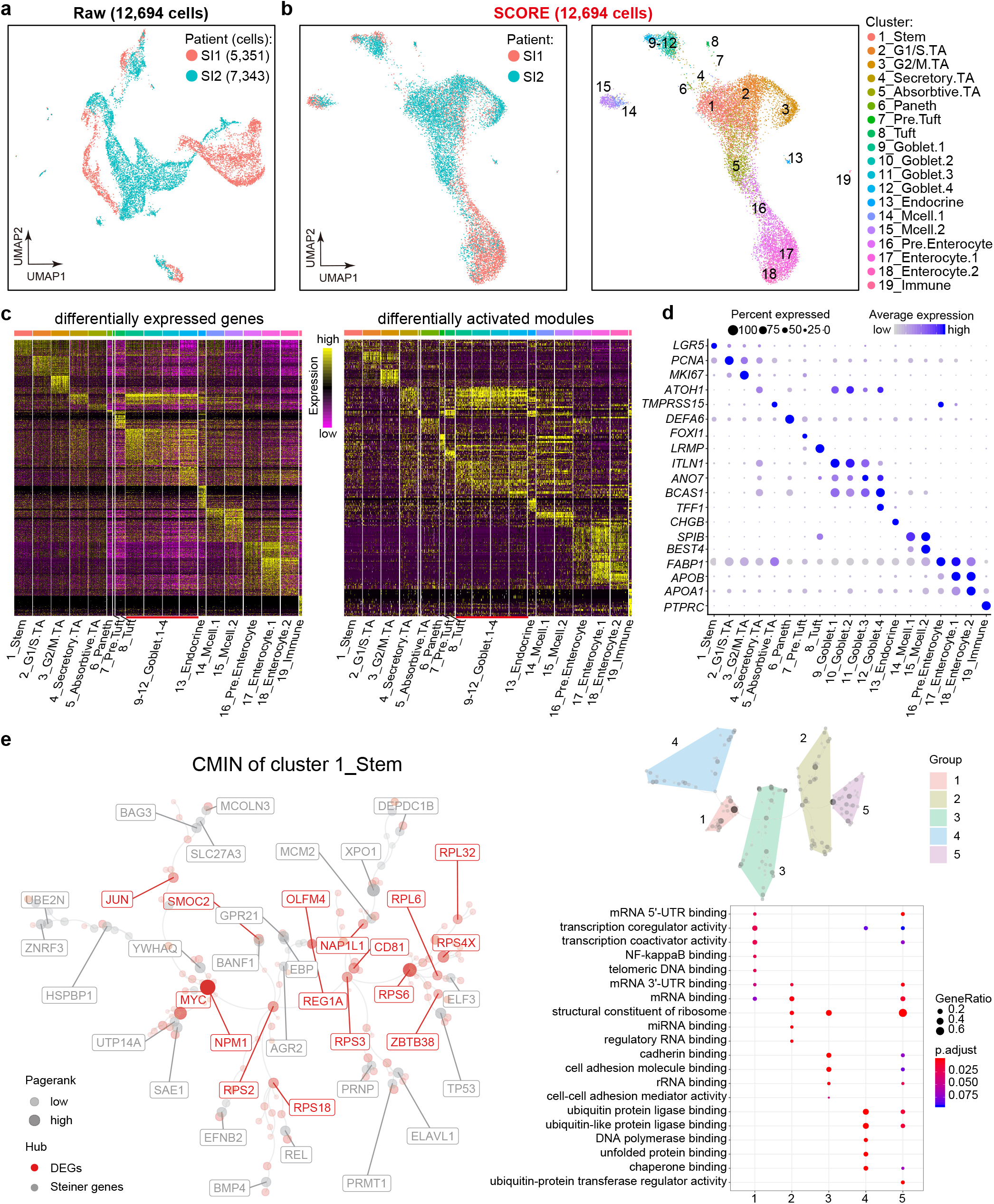
Expression landscape of human adult ileal dataset by SCORE. **a,** UMAP visualization of human adult ileal epithelium based on raw expression matrix within Seurat pipeline. Cells are colored by patient information. **b,** UMAP visualization based on SCORE. Cells are colored by patient information (left) and SCORE clusters (right). TA: transit-amplifying; Mcell: microfold cell. **c,** Heatmaps displaying DEGs (left) and DAMs (right) within each cluster. The color key from purple to yellow denotes low to high expression levels, respectively. **d,** Dotplot displaying the expression levels of representative marker genes of each cluster. Spot size denotes the percentage of cells expressing the gene within each cluster and colour intensity denotes the expression levels of the gene. **e,** CMIN of the epithelial stem/progenitor cells using the DEGs (left) and the related gene ontology terms of CMIN (right). The node sizes in CMIN represent the PageRank score of each gene and the genes of top 40 PageRank scores are displayed with their names in CMIN. The node colors denote the classification of genes as DEGs (red) and the connecting Steiner genes (gray) in CMIN. Spot size of gene ontology terms denotes the percentage of the related gene number and the color key denotes the adjusted p-values.

Importantly, with SCORE, we could construct the CMIN of a certain cluster to further explore the relationships and interactions of crucial genes (Fig. 4e and Supplementary Fig. 12b). Unlike traditional methods that focus on the differences in expression levels, CMIN ranks genes based on their topological importance in the optimal steiner tree. Therefore, the CMIN provides a simplified representation of original PPI network, and highlights the significance of non-marker interacting genes surpassing traditional marker gene analysis. Taking the cluster 1_Stem as an example, *MYC, CD81, OLFM4, JUN, REG1A, SMOC2, NPM1, NAP1L1* and several ribosome genes were inferred to be crucial for maintaining the cellular state of stem/progenitor cells (Fig. 4e). Moreover, the CMIN could be further divided into 5 gene groups based on their topological relationship, and the enriched gene ontology (GO) terms also supported the stem/progenitor identity of this cluster and improved the readability of the CMIN.

### 5. The heterogeneity within stem/progenitor cells, goblet cells and M cells

Despite the achievements in exploring the intestinal epithelium, a key question remains puzzling: do all intestinal stem cells have the equal differentiation potency? Using SCORE, we found that the stem/progenitor cells were heterogeneous and we identified 6 subgroups with the expression of specific marker genes, such as *TDGF1* (also known as Cripto-1), *GEM, FNBP1, ICOSLG,* and *DURAS3* (Fig. 5a,b). Cripto-1 plays an important role in early embryonic development, the formation and progression of several human tumor types^28^. Moreover, Cripto-1 can also interact with Wnt and Notch signaling pathways, which are crucial for the maintainence of intestineal stem cells^29^. Thus, the Cripto-1+ cells were impotant components of ileal epithelium, though their actual function remained to be explored.

**Fig. 5.**
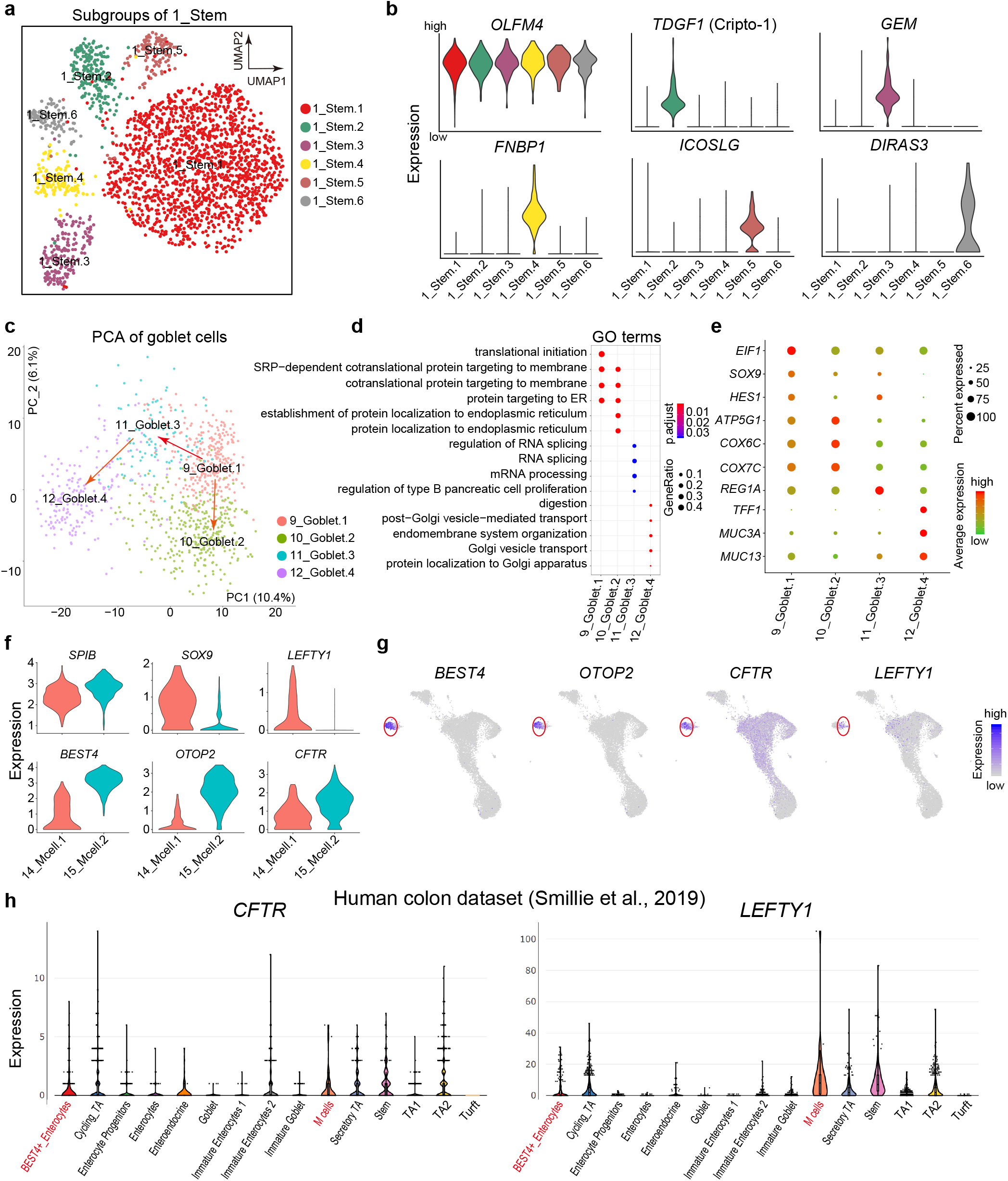
Subgroups of stem/progenitor cells, goblet cells and M cells. **a,** UMAP visualization of subgroups of cluster 1_Stem based on SCORE. Cells are colored by subgroups. **b,** Violinplots displaying the expression levels of representative marker genes of stem/progenitor subgroups. **c,** PCA plot displaying the subgroups of goblet cells. cells are colored by subgroups. **d,** Enriched GO terms using the DEGs of each goblet cell subgroup. Spot size denotes the percentage of the related gene number and the color key denotes the adjusted p-values. **e,** Dotplot displaying the expression levels of representative marker genes of goblet cell subgroups. Spot size denotes the percentage of cells expressing the gene within each cluster and colour intensity denotes their expression level. **f,** Violinplots displaying the expression levels of representative marker genes of two M cell subgroups. **g,** UMAP plots displaying the expression levels of representative marker genes across all the ileal cells. The color key denotes the expression levels. **h,** Violinplots displaying the expression levels of *CFTR* and *LEFTY1* in the human colon dataset published recently^32^.

As known, goblet cells secrete mucus to create a protective layer to the intestinal lumen^29^. We identified 4 subgroups of goblet cells with SCORE, namely, Goblet1-4, and they exhibited two varied differentiation routes: one is from Goblet1 to Goblet4 via Goblet3, while the other one is from Goblet1 to Goblet2 (Fig. 5c and Supplementary Fig. 13a). Of note, only one subgroup of goblet cells (8_Goblet4) is responsible for the secreation of mucus as they overrepresented *TFF1, MUC3A, MUC13;* while the other differentiation route, Goblet2, highly expressed genes related to respiratory electron transport, such as *ATP5G1*, *COX6C, COX7C.* (Fig. 5d,e).

Recently, a new cell type that distinctively expresses *BEST4* and *OTOP2* was found at the top of the colonic crypts, which can sense pH and transport salt, ions and metals^30^. In our dataset, we also detected this cell population and found that they were differentiated from the SPIB+ M cells (Fig. 5f and Supplementary Fig. 13b). Surprisingly, this *BEST4+/OTOP2+* M cells also highly expressed functional cystic fibrosis transmembrane conductance regulator *(CFTR)* (Fig. 5g). In human colon and mouse small intestine, *CFTR* is manly expressed in the crypt cells which helps these cells secrete fluid to flush the crypt lumen and remove contaminants^29^. In human ileal epithelium, the highest expression level of *CFTR* is restricted to the *BEST4+/OTOP2+* M cells, while this is not the situation in human colonic epithelium. Hence, *BEST4+/OTOP2+* M cells might play a different role in ileum compared with their colon counterpartners (Fig. 5h). Interestingly, the other M cell population highly expressed *LEFTY1,* which is also expressed in colonic M cells (Fig. 5g,h). *LEFTY1* encodes a secreted ligand that binds to Cripto-1 to antagonize Nodal signaling. Thus, these M cells might interact with the identified Cripto-1+ cells to regulate their differentiation or other cellular behaviors.

## Discussion

The close cooperation of different genes forms gene modules to fulfill specific cellular functions, and a certain cellular state or cell type can be well depicted by the activities of various gene modules^1^. As cellular states transit rapidly during organismal development and differentiation, how to simulate these dynamic processes and uncover the corresponding cell fates becomes a fundamental biology issue. In this study, we present a new computational method, SCORE, to infer this dynamic change and reveal cell development trajectory from the molecular network point of view. There are several advantages of SCORE compared with currently widely used methods.

Firstly, the hypothesis of SCORE is more biologically reasonable. Genes function in conjunction through molecular networks rather than in isolation. However, most published methods ignored the importance of molecular networks and just calculate the transcriptional similarity inferred from individual genes. A novel method called SCENIC utilizes transcription factor (TF) based networks to retrieve regulatory activity patterns, which can be used to annotate cellular states. However, SCENIC only measures the activity patterns of approximately 1500 TFs, and a recent evaluation showed that the sensitivity of SCENIC is only about 5% (about 75 TFs)^31^. Thus the resolution of SCENIC is limited, especially when analyzing highly heterogeneous single-cell data. In contrast, SCORE infers the molecular network from most of the expressed genes, and uses the network signatures rather than individual genes to define a certain cellular state, which highly improves the resolution and accuracy.

Secondly, SCORE can significantly reduce the false positive rates and yield more accurate results. SCORE uses the curated PPI network to correct the inferred gene-gene relationship, and all the interactions are literature-proved. As shown in Fig. 2, compared with other popular methods, SCORE possesses higher accuracy and robustness.

Thirdly, SCORE can overcome dropout effects and other technical variations to successfully integrate different datasets. One of the major drawbacks in scRNA-seq techniques is their high dropout rates. Since SCORE performs the analyses based on gene modules rather than individual genes, it can effectively reduce the unwanted dropout effects. In addition, PPI corrected correlation and AUC based module score are both able to eliminate the batch effects. As shown in Figs. 3 and 4, SCORE successfully integrated five human fetal datasets and one human adult ileal epithelium dataset, while other methods at least partially failed. Compared with current main-stream data integration methods, SCORE does not introduce artificial alternations to the raw gene expression matrix, nor require the input of exact batch ID for different datasets. The latter feature makes SCORE particularly useful to analyze time-series scRNA-seq datasets during development process, since the boundary between technical variations and biologically meaningful difference in various data collection time points is often blurred in such case. The subjective choice of batch ID for other data integration methods may omit the true temporal hetergeniety in gene expression.

Fourthly, in addition to the DEGs, SCORE can also identify the differentially activated gene modules (DAMs) and infer the gene relationship through the constructed CMIN for each cell fate (Fig. 4e). Moreover, due to the flexible framework of SCORE, it can be easily applied to deal with other networks, such as pathway network, metabolism network, etc. Finally, the molecular network decomposition step in SCORE workflow is realized with high efficiency and independent of data size. The major computational cost of SCORE lies in the quantification of module activities by AUCell, which scales linearly with cell numbers and has been easily paralleled in the R implementation. Thus, SCORE is highly scalable and is able to cater to the increasing size of scRNA-seq datasets nowadays.

However, as SCORE relies on the curated PPI network, it may not be applicable for the datasets of organisms without high-confidence molecular interaction information. Besides, when the available gene number is too low, the accuracy of SCORE may be impaired. Thus, we highly recommend researchers apply SCORE to analyze high-quality scRNA-seq datasets with greater sequencing depth.

In summary, with the high accuracy, robustness, and scalability, SCORE can help to explore the scRNA-seq datasets in a more biologically reasonable manner, and gain more insights into the complex biological systems.

## Supplementary figure and table legends

**Supplementary Fig. 1.** The performance of SCORE on the cell-line benchmarking dataset was compared with the direct analysis on raw expression matrix within Seurat pipeline (denoted as Raw) and other gene regulatory network (GRN) based methods (i.e. CSN and SCENIC with continuous/binary features). The number of input highlight variable genes is set as 14,000, and the colors in t-SNE plot denote cell lines identity collected in the experiments.

**Supplementary Fig. 2.** The performance of SCORE on the cell-line benchmarking dataset was compared with the direct analysis on raw expression matrix within Seurat pipeline (denoted as Raw) and other gene regulatory network (GRN) based methods (i.e. CSN and SCENIC with continuous/binary features). The number of input highlight variable genes is set as 11,000, and the colors in t-SNE plot denote cell lines identity collected in the experiments.

**Supplementary Fig. 3.** The performance of SCORE on the cell-line benchmarking dataset was compared with the direct analysis on raw expression matrix within Seurat pipeline (denoted as Raw) and other gene regulatory network (GRN) based methods (i.e. CSN and SCENIC with continuous/binary features). The number of input highlight variable genes is set as 8,000, and the colors in t-SNE plot denote cell lines identity collected in the experiments.

**Supplementary Fig. 4.** The performance of SCORE on the cell-line benchmarking dataset was compared with the direct analysis on raw expression matrix within Seurat pipeline (denoted as Raw) and other gene regulatory network (GRN) based methods (i.e. CSN and SCENIC with continuous/binary features). The number of input highlight variable genes is set as 2,000, and the colors in t-SNE plot denote cell lines identity collected in the experiments.

**Supplementary Fig. 5.** Comparison of marker gene analysis based on different data integration results by SCORE and other methods, with the top 10 marker genes displayed for each cluster. The columns in the matrices represent cells, and the rows represent genes. The color key from blue to red denotes low to high expression levels, respectively. The incorrectly clustered cells, which do not significantly express marker genes of the assigned cluster, are marked by the black squares.

**Supplementary Fig. 6. Performance comparison between standard Seurat pipeline (denoted as Raw) and SCORE on human fetal kidney dataset.**

**a,** UMAP visualization of cell clusters of kidney dataset based on standard Seurat pipeline (denoted as Raw). Cells are colored by cell types (left) and developmental weeks (right).

**b,** Feature plots displaying the expression levels of representative marker genes. The color key from grey to red denotes low to high expression levels, respectively.

**c,** UMAP visualization of cell clusters of kidney dataset based on SCORE. Cells are colored by cell types (left) and developmental weeks (right).

**d,** Feature plots displaying the expression levels of representative marker genes.

**e,** Schematic diagram of renal tubule.

**Supplementary Fig. 7. Performance comparison between standard Seurat pipeline (denoted as Raw) and SCORE on human fetal heart dataset.**

**a,** UMAP visualization of cell clusters of heart dataset based on standard Seurat pipeline (denoted as Raw). Cells are colored by cell types (left) and developmental weeks (right).

**b,** Feature plots displaying the expression levels of representative marker genes. The color key from grey to red denotes low to high expression levels, respectively.

**c,** UMAP visualization of cell clusters of heart dataset based on SCORE. Cells are colored by cell types (left) and developmental weeks (right).

**d,** Feature plots displaying the expression levels of representative marker genes.

**e,** UMAP plots of subgroups of cardiomyocytes (CMs) based on SCORE (left). Cells are colored by subgroups. Feature plots displaying the expression levels of representative marker genes of subgroups (right).

**Supplementary Fig. 8. Performance comparison between standard Seurat pipeline (denoted as Raw) and SCORE on human fetal gonad dataset.**

**a,** UMAP visualization of cell clusters of fetal gonad dataset based on standard Seurat pipeline (denoted as Raw). Cells are colored by cell types (left) and developmental weeks (right).

**b,** Feature plots displaying the expression levels of representative marker genes. The color key from grey to red denotes low to high expression levels, respectively.

**c,** UMAP visualization of cell clusters of fetal gonad dataset based on SCORE. Cells are colored by cell types (left) and developmental weeks (right).

**d,** Feature plots displaying the expression levels of representative marker genes.

**Supplementary Fig. 9. Performance comparison between standard Seurat pipeline (denoted as Raw) and SCORE on human fetal cerebral cortex dataset.**

**a,** UMAP visualization of cell clusters of cerebral cortex dataset based on standard Seurat pipeline (denoted as Raw). Cells are colored by cell types (left) and developmental weeks (right).

**b,** Feature plots displaying the expression levels of representative marker genes. The color key from grey to red denotes low to high expression levels, respectively.

**c,** UMAP visualization of cell clusters of cerebral cortex dataset based on SCORE. Cells are colored by cell types (left) and developmental weeks (right).

**d,** Feature plots displaying the expression levels of representative marker genes.

**Supplementary Fig. 10. Performance comparison between standard Seurat pipeline (denoted as Raw) and SCORE on human fetal prefrontal cortex dataset.**

**a,** UMAP visualization of cell clusters of prefrontal cortex dataset based on standard

Seurat pipeline (denoted as Raw). Cells are colored by cell types (left) and developmental weeks (right).

**b,** Feature plots displaying the expression levels of representative marker genes. The color key from grey to red denotes low to high expression levels, respectively.

**c,** UMAP visualization of cell clusters of prefrontal cortex dataset based on SCORE. Cells are colored by cell types (left) and developmental weeks (right).

**d,** Feature plots displaying the expression levels of representative marker genes.

**Supplementary Fig. 11. Quality control of two human adult ileal samples.**

**a,** Histological sections of the two sampled human adult ileal tissues.

**b,** 10X dataset quality control information.

**Supplementary Fig. 12. Characterizations of the human adult ileal cell types.**

**a,** Boxplot displaying the differentiation potency inferred by the SCENT algorithm.

**b,** CMINs of representative clusters using the DEGs. The node sizes in CMIN represent the PageRank score of each gene and the genes of top 20 PageRank scores are displayed with their names in CMIN. The node colors denote the classification of genes as DEGs (red) and the connecting Steiner genes (gray) in CMIN.

**Supplementary Fig. 13. Expression patterns of goblet cell and microfold cell subgroups.**

**a,** Heatmap displaying the DEGs within each goblet cell subgroup. The top 10 marker genes were listed on the right. The color key from purple to yellow denotes low to high expression levels, respectively.

**b,** Heatmap displaying the DEGs within each microfold cell subgroup. The top 10 marker genes were listed on the right.

**Supplementary Table1. Cell information and DEGs of human adult ileal dataset.**

## Methods

### Overview of SCORE

For the convenience of users, SCORE is seamlessly compatible with the Seurat pipeline. The input for SCORE workflow includes a single-cell gene expression matrix and a PPI network. Both the cells and genes in the expression matrix could be pre-filtered by users, while it is highly recommended that enough number of genes (>5000) should be retained to achieve more robust analysis of SCORE, as shown via the gold-standard cell line dataset (see main text and SI). The nodes of PPI network should overlap considerably with the gene names in the expression matrix, and the edges of the network shall represent the corresponding molecular interactions with relatively high confidence. In the R implementation of SCORE, the procedure supports the automatic download of PPI from public database such as Biogrid and STRING, which is recommended in the standard workflow. The output of SCORE is a cell-module matrix representing the activity of individual dynamic module within each cell, which can be utilized for downstream visualization, clustering and cell lineage analysis.

### Dissection of dynamic molecular networks

To simultaneously reduce the false-positive interactions from correlation inference, and prune data-irrelevant interactions of PPI network, SCORE constructs a weighted molecular interaction network (WMIN) *G*(*V, E, W*) by combining the data-driven and knowledge-based approaches. The vertex set *V* of the network only consists of the genes in the input expression matrix, and the edge set *E* obtained from the input PPI network. The weight *W_ij_* on the edge *E_ij_* is the Pearson correlation coelficients between the corresponding nodes *i* and *j* calculated from the single-cell gene expression matrix. To improve the interpretability of molecular networks and highlight the co-expression features, by default we only keep the edges with positive weight in the WMIN.

To achieve fast and robust identification of dynamic molecular networks, SCORE utilizes the consensus detection of the weighted network community through random walk approach. Given the weighted network *G*(*V, E, W*), a random walk on the network is naturally induced, whose transition probability matrix (TPM) *P* is defined by

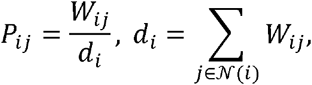

Where *N*(*i*) denotes the neighbors of the node *i.* SCORE constructs an ensemble of random walks with different step lengths on the WMIN, with the TPMs *P^l_1_^,P^l_2_^, …,P^1_R_^*, where the power of matrix *l_m_* denotes the length of time step. The walktrap algorithm is applied to detect network community for each random walk, respectively. The algorithm partitions the network in terms of the distance induced by the random walk, based on the intuition that the random walker will be “trapped” in the closely-connected sub-networks (termed as modules). All modules with molecule number larger than 3 in each run will be kept as the final dynamic modules, resulting in the module set 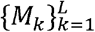. It would be possible that certain modules occur repeatedly in different runs of walktrap algorithm, indicating their stability to form a closely-connected community. SCORE strengthens the weight of such modules automatically in the subsequent analyses, by restoring all the modules without deletion of repeated items.

### Quantification of the module activity

For each detected module, SCORE utilizes AUCell to quantify its activation level within each individual cell. Given cell *x,* genes are ranked in descending order according to their expression level in *x.* By default, the z-score is adopted in SCORE to rank the genes in order to remove the effect of scaling. The recovery curve (ROC) for module *M_k_* is then derived by counting the top ranked genes enriched in *M_k_.* The activity measure *A_k_(x)* of module *M_k_* in cell *x* (consists of the final output matrix of SCORE) is defined as the area under the curve (AUC) for the top ranked genes. Intuitively, modules with higher activity in the biochemical process tend to possess the high-ranking gene expression level, therefore associate with higher activity measure. The AUCell procedure is independent of gene expression unit or normalization method, therefore achieving the effective removal of batch effect in the single-cell experiments.

### Downstream Analysis

The obtained module activity *A_k_*(*x*) matrix from SCORE (whose rows represent modules and columns represent cells) can replace the raw gene expression matrix as the input for downstream analysis, such as dimension reduction, clustering and lineage inference. As shown in various datasets of the main text, the downstream analysis based on SCORE module activity features outperforms the raw expression matrix, in terms of clustering accuracy, development lineage trend, and removal of experimental batch effects. Hence, the workflow of SCORE can be understood as the extraction of biologically meaningful and robust features, guided by molecular interactions in the single-cell transcriptome data.

For the convenience of downstream analysis, the R implementation of SCORE is deeply fused with the workflow of Seurat v3.0 package. The input expression matrix to SCORE can be a Seurat object with RNA assay, and the output module activity features are returned as the Net assay in the same Seurat object. Users may conveniently conduct dimension reduction and cell-clustering based on the Net assay, and perform marker gene analysis based on the RNA assay, by switching the default assay of Seurat object.

### Construction of Cell State-Specific Characteristic Molecular Interaction Network (CMIN)

To annotate a certain cell state, beyond the marker genes, **SCORE** constructs the CMIN with the concept of Steiner Tree in graph theory. Given a graph *G =* (*V, E*) and a subset of vertices *T* ⊂ *V* (called terminal vertices), a Steiner tree *S* ⊂ *G* is a connected tree that spans through the given terminal vertices *T*. The Steiner tree *S* may contain vertices not presented *in T,* known as the Steiner vertices, serving as the interchange node to connect the vertices *in T*. In molecular interaction network, the marker genes of a certain cell state are typically selected as the terminal vertices, and the mediating Steiner vertices, although not necessarily differentially expressed, are supposed to play important roles in formulating the specific cell state through molecular interactions. Therefore, the Steiner tree provides a simplified representation of original PPI network, and highlights the significance of non-marker interacting genes surpassing traditional marker gene analysis.

In the downstream analysis of **SCORE**, given the specific cell cluster identified from module activation features, we first construct the set of terminal genes *T\** by detecting the marker genes from two different levels,

1) Union of all genes in the **SCORE**-extracted modules that are differentially activated in the cluster, denoted as differentially activated module genes (DAMGs);

2) Individual genes that are significantly up-regulated in the cluster, denoted as differentially expressed genes (DEGs).

While the DEGs are commonly referred as the “markers” of cellular states, the DAMGs also represent the key molecular interaction modules to mark the cell cluster in the network resolution.

Next, to infer the CMINs that possibly formulate the cellular states rather than the genes solely marking the cellular states, we propose to calculate a Steiner tree *S\** that spans the terminal gene set *T*\* with some optimal property, defined on the constructed WMIN *G*(*V,E,W*) by **SCORE** in the first step of workflow. We require that the separate DAMGs or DEGs in *S\** are linked by the most relevant genes, as well as through the most likely interaction path derived from the dataset. To this end, we define the distance *D_ij_*, on the edge *E_ij_* of WMIN by *D_ij_, = l/(W_ij_* + ε), where the small number ε is added to avoid zero in dominator. Highly correlated gene pairs in PPI tend to possess much closer distance from the definition. Then S* can be optimized as the Steiner tree with the least sum of edge distances, which can be tackled efficiently by the greedy algorithm.

For a better visualization of CMIN, in R implementation of **SCORE** we mark DAMGs, DEGs, and Steiner connecting genes with different colors, and also use the PageRank algorithm to measure the topological importance of the genes in optimal Steiner tree *S\** as shown by the size of the nodes. We can also provide any two genesets to construct the CMIN.

### Settings in the benchmarking gold-standard dataset

The gene expression matrix was processed as fragments per kilobase per million reads (FPKM) as in the original literature. To test the robustness of different methods, we first adopted the vst method in Seurat v3.0 package to select five groups of highly variable genes (with the number of genes 2,000, 5,000, 8,000, 11,000, and 14,000, respectively). We performed SCORE, as well as two other network or biological information based methods, CSN and SCENIC to further compress and extract the features, respectively, from the groups of highly variable genes (HVGs).

For SCORE, we downloaded the *Homo sapiens* PPI network (version 3.5.173) from the BioGRID database. The top ranked genes included in the calculation of AUC values varied with the sizes of input HVGs, with 250 and 200 for 2,000 and 5,000 variable genes, respectively, and 400 in other cases. The parameters in implementing CSN and SCENIC were chosen with default values. The SCENIC yields both continuous and binary features as the outputs.

The running time comparison was conducted on the 2.50GHz Xeon E5-2680 machine with 128G RAM, 12 cores and Linux OS. For SCORE and SCENIC, the CPU core number was set as 10. CSN was automatically paralleled with MATLAB 2019b. The wall time of implementing each procedure were recorded as the running time in the main text.

In the downstream clustering analysis, three methods, SNN, SIMLR and SC3 were performed on the extracted features by different methods, and the adjusted rand index (ARI) as well as the similarity matrix of SC3 were used to evaluate the accuracy and the effect of batch removal. The direct analysis on raw expression matrix with selected HVGs was also performed for the comparison. The true labels were the seven collected cell lines identity (H1, GM12878, A549, HCT116, H1437, K562 and IMR90) without batch information. For SNN, we tuned the resolution parameter to obtain the optimal ARI value. As to SC3 and SIMLR, we set the number of clusters to 7. The t-SNE plot was produced based on the top 10 principal components of the extracted features.

### Human fetal datasets

To evaluate the integration performance of SCORE, five human fetal datasets were collected from our previously published studies, including fetal gonads (overies and testis) (GEO number: GSE86146), heart (GEO number: GSE106118), kidney (GEO number: GSE109488), prefrontal cortex (PFC) (GEO number: GSE104276), and cerebral cortex (GEO number: GSE103723), spanning from 4 to 26 weeks of fetal development. Importantly, we re-organized the five datasets using uniform pipeline and format, which provided a rich and convenient resource for studying human fetal development (https://github.com/zorrodong/HECA). In brief, barcode and UMI information were extracted by UMI-tools form raw reads^33^. After discarding the poly A bases, TSO sequences and low-quality sequences, the clean reads were mapped to GRCh38 reference using STAR aligner^34^. We used featureCounts^35^ to annotate the mapped reads and quantified the UMI counts through UMI-tools. We provided the pipeline for users (https://github.com/zorrodong/HECA).

To analyze the human fetal datasets, we first discard cells with gene number below 1,000 and UMI counts below 10,000. HVGs were chosen using Seurat (mean >= 0.1, dispersion >= 0.1), and about 8,000 HVGs were selected for SCORE to perform the evaluation. Two batch effect correction methods CCA and Harmony were used with the recommended parameters. 30-50 reduced dimensions were used to perform UMAP analysis and clustering using the graph-based method in Seurat. To accelerate the speed, we used genesorteR to conduct the differential expression analysis.

### Human adult ileal crypt dataset

This study was approved by the Ethics Committee of Peking University Third Hospital (License No. M2016170). All patients had signed written informed consent. Ileal crypt samples were collected from 2 right-sided colon cancer patients immediately after surgical resection. We dissociate the samples into single cells and constructed the libraries using 10x Genomics (3’ Library, Kit v3). Libraries were sequenced using Illumina HiSeq 4000 platform with 150-bp paired-end reads.

We used Cellranger v3.1.0 (10X Genomics) to deal with the raw reads and quantify the expression level. Next, the UMI count matrix were analyzed using Seurat pipeline. We discarded cells with gene number below 1,000, UMI counts below 1,000, and mitochondrial percentage above 30%. 8,000 HVGs were chosen for SCORE analysis using FindVariableFeatures(nfeatures = 8000,selection.method = “vst”). The overall dimensionality reduction and clustering were performed using all the obtained modules.

### Data availability

Human adult small intestine dataset are deposited in the GEO. SCORE is freely available in https://github.com/wycwycpku/RSCORE. The five human fetal datasets are available in https://github.com/zorrodong/HECA.

## Acknowledgements

This project was supported by grants from the National Natural Science Foundation of China (31625018 and 81521002 to F.T., 11825102 and 11421101 to T.L.).

## Author contributions

F.T., T.L., W.F., J.D., P.Z. conceived the project; W.W., X.Z., Y.G., L.W., performed the experiments; J.D., P.Z., Y.W., Y.C., H.X., J.L., J.Y., X.N.Z. conducted the bioinformatics analyses; J.D., P.Z., T.Li, F.T., wrote the manuscript with the help of all the authors.

